# A neutral process of genome reduction in marine bacterioplankton

**DOI:** 10.1101/2024.02.04.578831

**Authors:** Xiaojun Wang, Mei Xie, Kaitlyn Elizabeth Yee Kei Ho, Ying Sun, Xiao Chu, Shuangfei Zhang, Victoria Ringel, Hui Wang, Xiao-Hua Zhang, Zongze Shao, Yanlin Zhao, Thorsten Brinkhoff, Jörn Petersen, Irene Wagner-Döbler, Haiwei Luo

## Abstract

Marine bacterioplankton communities are dominated by cells equipped with small genomes. Streamlining selection has been accepted as the main force driving their genome reduction. Here, we report that a neutral evolutionary mechanism governs genome reduction in the Roseobacter group that represents 5-20% of the bacterioplankton cells in coastal waters. Using representative strains that fall into three genome size groups (2-3, 3-4, and 4-5 Mbp), we measured their genomic mutation rates (μ) through long-term mutation accumulation experiments followed by genome sequencing the resulting 437 mutant lines. We further calculated their effective population sizes (*N_e_*) based on μ and the neutral genetic diversity of the studied species, the latter estimated based on multiple genome sequences of natural isolates collected from global oceans with their population structure considered. A surprising finding is that *N_e_* scales positively with genome size, which is the opposite of the expectation from the streamlining selection theory. As the strength of random genetic drift is the inverse of *N_e_*, this result instead suggests drift as the primary driver of genome reduction. Additionally, we report a negative scaling between μ and genome size, which is the first experimental evidence for the long-lasting hypothesis that mutation rate increases play a part in marine bacterial genome reduction. As μ scales inversely with *N_e_*, genetic drift appears to be the ultimate cause of genome reduction in these Roseobacters. Our finding discounts, but is insufficient to reject, the streamlining theory because streamlining process is expected to be more effective in oligotrophic open ocean waters.

## Introduction

Bacterioplankton in the surface ocean is dominated by genome-reduced lineages, which are characterized by an average genome size of less than 2 Mbp ^1–3^. Among these, the most abundant genome-reduced lineages are the SAR11 clade (Pelagibacteriales) of Alphaproteobacteria, the SAR86 clade of Gammaproteobacteria, and the genus *Prochlorococcus* of Cyanobacteria ^4^. Streamlining selection is believed to drive the genome reduction of these free-living bacteria ^5,6^. According to this theory, selection favors smaller genome size in nutrient-limited surface oceans because i) removal of non-essential DNA provides a metabolic advantage for bacteria, and ii) a concomitant increased surface-to-volume ratio (correlated with a decreased genome size) promotes nutrient uptake through enhanced diffusive delivery of nutrients to cell surface ^6–8^. The key concept of streamlining selection theory hinges on the effective population size (*N_e_*), which describes how many individuals in a bacterial population are contributing to the observed neutral genetic diversity of the population ^9^. The underlying hypothesis is that free-living marine bacteria with reduced genomes have larger *N_e_*than their relatives carrying larger genomes, thereby increasing the efficiency of natural selection acting to eliminate superfluous genomic DNA in oligotrophic environments. By comparing the *N_e_* of related lineages with very different genome sizes, we can start to assess the definitive role of selection to downsize the genomes of free-living bacterioplankton cells in surface oceans.

However, this hypothesis has never been tested experimentally. The *N_e_* can be calculated if π_S_ (the neutral genetic diversity of a population) and µ (unbiased genomic mutation rate) are available according to the equation π_S_ = 2 x N xµ ^10^. Here, π_s_ is approximated by the diversity at four-fold degenerate sites of a population whose members recombine more frequently than those involving other populations. As a gold standard, µ is measured using a long-term mutation accumulation (MA) experiment, which requires the experimental populations to repeatedly go through single-colony bottlenecks to eliminate the effect of natural selection ^11^. This is commonly achieved by growing bacterial cells to single colonies in many parallel lines and propagating them for hundreds of generations. By whole genome sequencing (WGS) the mutant lines and comparing the mutant genomes with the ancestor’s genome, the unbiased genomic mutation rate and spectrum can be calculated. However, genome-reduced bacterioplankton lineages inhabiting surface oceans typically do not grow on solid media or do not form single colonies on solid media, thus posing a significant challenge for their genomic mutation rate measurements and *N_e_*estimation. A genome-reduced member of *Prochlorococcus* in the high-light-adapted clade II (*Prochlorococcus marinus* AS9601, ∼1.7 Mbp) is the only genome-reduced surface ocean bacterioplankton lineage that was subjected to MA/WGS procedure ^12^.

The Roseobacter group (Roseobacteraceae, Alphaproteobacteria) is globally abundant, comprising up to 20% of the bacterial cells in coastal waters and 5% in the open ocean ^13–15^. Cultured members of the Roseobacter group exhibit a wide range of genome size spanning from 2.5-2.6 Mbp (CHUG, NAC11-7) to 6.5-8.1 Mbp (*Salipiger*, *Poseidonocella*) (with the 25%, 50%, and 75% percentile being 3.83, 4.36, and 4.69 Mbp, respectively, Fig. S1). Most cultured lineages thrive on nutrient-rich solid media, but they are generally not representative in the oligotrophic pelagic ocean environments ^16^. Early community structure analyses based on 16S rRNA gene revealed that marine Roseobacter communities are dominated by a few uncultivated lineages ^17^, whose genomes are becoming increasingly available through sequencing novel cultured members and sequencing uncultured members by single-cell genomics and metagenomic binning. Classical examples are DC5-80-3 (also named RCA or *Planktomarina*) ^18–21^, CHAB-I-5 ^15,22^, ChesI-A/B (also named SAG-O19) ^23^, NAC11-7 ^24^, and more recently CHUG ^25,26^. These lineages constitute a polyphyletic group in the phylogenomic tree but form a tight pelagic Roseobacter cluster (PRC) in a dendrogram clustered from the presence and absence of gene families, reflecting their convergent evolution towards shared genome content and reduced genome sizes ^22,26^.

Unlike other PRC lineages, some CHUG isolates grow in single colonies and can be stably propagated on agar plates, thus rendering themselves appropriate for genomic mutation rate measurement through the classical MA/WGS procedure. Since other members of the Roseobacter group that carry larger and variable genomes and co-inhabit surface oceans are more readily available for µ determination, CHUG and its Roseobacter relatives commonly found in surface oceans create a unique opportunity to test the streamlining selection hypothesis.

Here, we report the µ of CHUG to be (7.86 ± 5.31) × 10^-10^ base substitutions per nucleotide site per cell division, which, to date, represents the free-living bacterial species with the highest mutation rate measured by MA/WGS. Additionally, we report the *N_e_* of CHUG to be 1.78 X 10^7^, which, together with *Prochlorococcus* high light-adapted clade II (1.68 X 10^7^) ^12^, represent the free-living bacterial species with the smallest *N_e_*. We also determined µ and *N_e_*of two other Roseobacter species, namely *Sulfitobacter pontiacus* and *Dinoroseobacter shibae*, and included the published *Ruegeria pomeroyi* data ^20^ for comparison. These four Roseobacter species are all commonly found in surface ocean habitats but have overall very different genome sizes (2.6, 3.5, 4.4, 4.6 Mbp). Remarkably, the assayed Roseobacter species that carry lower genome sizes have higher µ and smaller *N_e_*. These scaling relationships imply that both genetic drift (i.e., the strength of drift is the inverse of *N_e_*) and mutation rate increases play important roles in genome reduction of this bacterioplankton group. That *N_e_* scales inversely with µ further implies that genetic drift is the ultimate force that governs genome reduction. This is the first experimental test of the prevailing streamlining selection theory and the finding is surprising because it reverses the trend predicted by the streamlining theory in which streamlined marine bacterioplankton lineages should have very large *N_e_* such that genetic drift is negligible.

## Results

### Genome sizes of the Roseobacter group members from surface oceans scale negatively with their genomic mutation rates

Here, we report the genomic mutation rate (μ) of three important members of the Roseobacter group: CHUG, *Sulfitobacter pontiacus*, and *Dinoroseobacter shibae*. Along with μ of *Ruegeria pomeroyi* we published earlier ^20^, we have four representative Roseobacter lineages commonly found in surface ocean habitats and varying substantively in genome sizes (2.6, 3.5, 4.4 and 4.6 Mbp, respectively). For CHUG, a total of 200 mutation accumulation (MA) lines were initiated from a single ancestral cell of strain HKCCA1288, 192 of which survived after 64 transfers with each line undergoing 1,472 cell divisions (corrected with death rate) and 180 of which had over 50x coverage in WGS. Mutations were accumulated in 172 out of the 180 MA lines, and a total of 596 base-pair substitution mutations (BPSs), 121 deletions and 29 insertions were identified (Table S1).

In general, mutations are randomly distributed along the genomic regions of CHUG, though clustered mutations were identified in two ribosomal RNA (rRNA) genes and 17 protein-coding genes across multiple MA lines (Fig. 1A). In the former, 25 and 15 BPSs are clustered within 23S rRNA and 16S rRNA genes, respectively. Among these, 28 BPSs are contributed by a single MA line and the remaining 12 are distributed across another 12 MA lines (Table S2). We validated a randomly chosen BPS located in the 23S rRNA gene using PCR (Table S2). For the latter, 43 BPSs, three deletions, and 72 insertions fell into 17 genes across 72 MA lines, which represents a significant excess of mutations (bootstrap test; *p* < 0.05 for each gene, Fig. 1A).

**Fig. 1.**
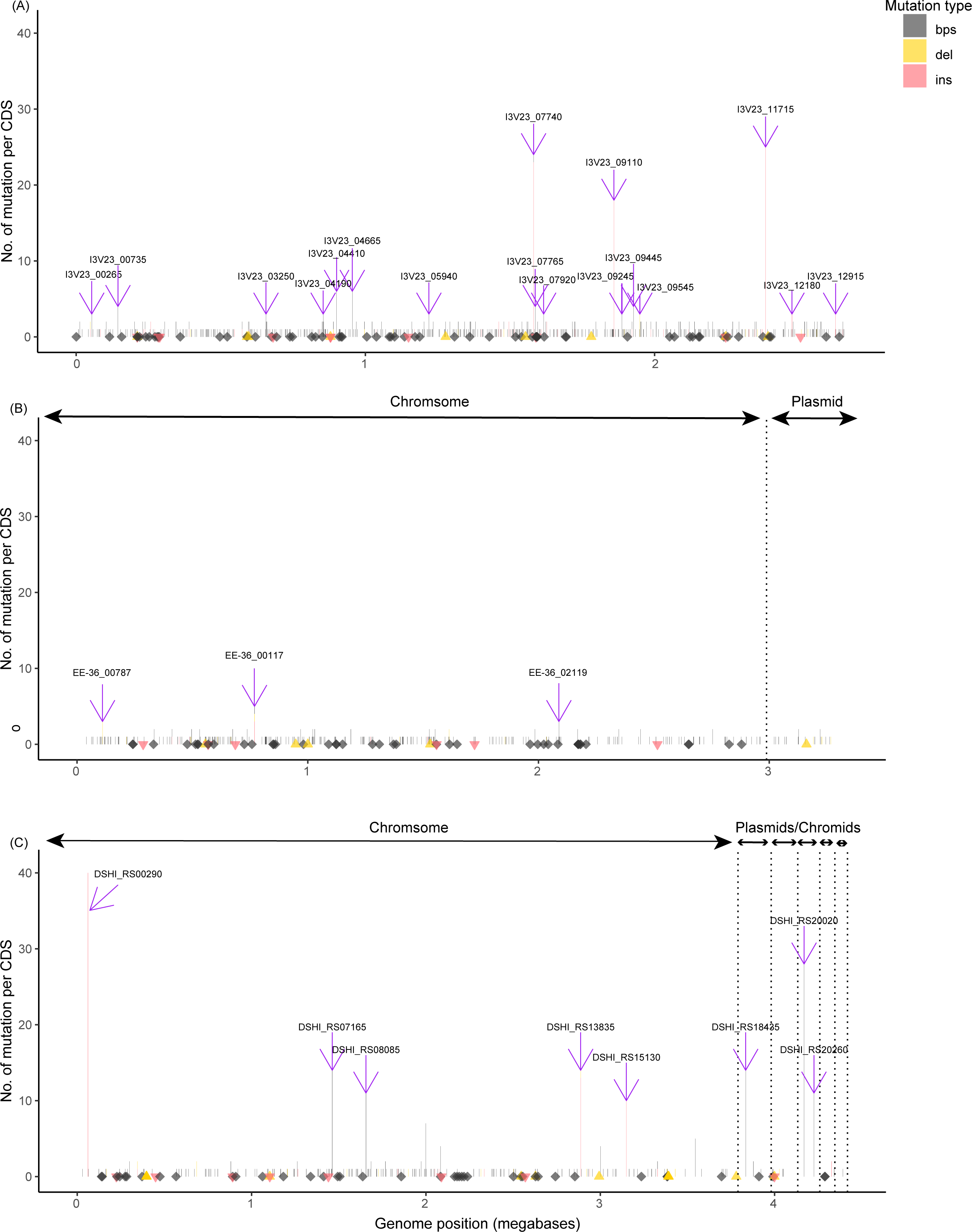
Distribution of genomic mutations in three Roseobacter species determined in this study by mutation accumulation (MA) experiments followed by whole genome sequencing of 437 MA lines in total. (A) Base-substitution mutations and insertion/deletion mutations across the whole genome of CHUG HKCCA1288 determined by 180 MA lines. The height of each bar represents the number of base substitutions (black), insertions (red) and deletions (yellow) across all MA lines within each protein-coding gene. Black diamonds and red triangles denote base substitutions and insertions that occurred on the remaining genomic regions (intergenic regions and non-protein-coding genes), respectively; both diamonds and triangles are shown with transparence, thus genomic regions with more mutations show deeper color than those with less mutations. The genomic position of insertion/deletion mutation refers to the position of the first mutated site. The locus tag of the 17 genes with statistical enrichment of mutations is shown in purple arrows. (B) Base-substitution mutations and insertion/deletion mutations across the whole genome of *Sulfitobacter pontiacus* EE-36 determined by 58 MA lines. Different types of mutations are illustrated with the same symbols as those in (A). The locus tag of the three genes with statistical enrichment of mutations is shown in purple arrows. Chromosome and plasmid are separated with the vertical dash line. (C) Base-substitution mutations and insertion/deletion mutations across the whole genome of *Dinoroseobacter shibae* DFL-12 determined by 149 MA lines. Different types of mutations are illustrated with the same symbols as those in (A). The locus tag of the eight genes with statistical enrichment of mutations is shown in purple arrows. Chromosome, chromids, and plasmids are separated with the vertical dash line.

Most of the insertional mutations (66 of the 72) fall into three genes (I3V23_07740, I3V23_09110, and I3V23_11715), and most of them (60 of the 66) are frameshift mutations (Table S2), potentially leading to pseudogenization. Gene I3V23_07740 encodes the SLAC1 anion channel family protein linked to tellurite resistance, I3V23_09110 produces the DeoR/GlpR transcriptional regulator associated with fructose and glucose metabolism, and I3V23_11715 codes for the sarcosine oxidase subunit beta family protein (Table S2). Mutation clustering is a common phenomenon in mutation rate determination with the MA/WGS strategy ^12,27–29^, and it may be either caused by mutational hotspots or a result of positive selection as they likely increase fitness under experimental conditions. In the case of CHUG, frameshift mutations are enriched in the three genes, tentatively suggesting that deleting these genes likely confer benefits to the bacteria during the MA process. Thus, BPSs occurring in the above genes or intergenic region were excluded when calculating the genomic base-substitution mutation rate, and the remaining 505 BPSs translate to a μ of (7.31 ± 4.92) × 10^-10^ (95% confidence interval [CI]: 6.69 × 10^-10^ – 7.97 × 10^-10^) base substitutions per site per cell division.

If the mutations clustered in the above genomic regions are indeed under selection, they may hitchhike mutations in linked genomic regions ^30,31^, thus inflating the mutation rate estimate. They may also have epistatic interactions with mutations in other genomic regions ^32,33^, likely delaying the fixation of other mutations and thus reducing the mutation rate estimate. To eliminate any such confounding factors in mutation rate estimation, we discarded the 83 MA lines with mutations accumulated in any of the 17 genes or the two ribosomal RNA genes with excess mutations. The remaining 97 MA lines accumulated 270 BPSs, 19 deletions and 28 insertions (Table S1), translating to a μ of (7.14 ± 4.82) × 10^-10^ (95% CI: 6.32 × 10^-10^ – 8.05 × 10^-10^), which is not significantly different from the mutation rate (7.51 ± 5.03) × 10^-10^ (95% CI: 6.60 × 10^-10^ – 8.52 × 10^-10^) derived from the remaining 83 MA lines mutated at aforementioned genes but with these mutations excluded from the calculation (Wilcoxon–Mann–Whitney test, *p* = 0.930).

Additional analyses were performed to evaluate the effect of selective pressure in the 97 MA lines where mutations are randomly distributed across the entire genomes. We found the ratio of accumulated mutations in protein-coding sites to those in intergenic sites (233 vs 1) is significantly larger than the ratio of the number of protein-coding sites to intergenic sites (2,432,502 vs 214,229) (Fisher’s exact test, *p* < 0.001). A similar pattern has also been reported in MA experiments with other bacteria ^34,35^, and the increased mutation rate in coding regions may be linked to transcription-induced mutations, considering that coding regions are transcribed whereas most intergenic regions are not ^36,37^. In terms of the protein-coding sites, the ratio of accumulated nonsynonymous to synonymous mutations (171 vs 62) does not differ significantly from that of nonsynonymous to synonymous sites (1,766,993 vs 665,509) (χ^2^ test, *p* = 0.85), supporting that selection did not play an important role during the MA process.

For the *Sulfitobacter* and *Dinoroseobacter*, MA experiments were conducted using two strains, *S. pontiacus* EE-36 (=DSM 11700) and *D. shibae* DFL-12 (=DSM 16493^T^), and the same mutation calling method was implemented to calculate the spontaneous mutation rates. For EE-36, an analysis of 58 MA lines with high-quality reads revealed 284 BPSs, 22 deletions and 14 insertions (Table S3&S4 and SI Results). Among the 51 MA lines without the potential effect of the three mutation-enriched genes found here (Fig. 1C and SI Text 1.1), 243 BPSs, 11 insertions and 18 deletions (Table S3) were kept, and the 243 BPSs translate to a μ of (3.57 ± 1.89) × 10^-10^ (95% CI: 3.13 × 10^-10^ – 4.03 × 10^-10^). No selection was detected in the MA experiment of EE-36 (SI Text 1.1). For DFL-12, 149 MA lines with high quality reads yield 296 BPSs, 34 deletions and 82 insertions (Table S5&S6). After ignoring 87 MA lines potentially affected by the eight genes with an excess of mutations, the remaining 62 MA lines with 80 BPS lead to a μ of (1.81 ± 2.25) × 10^-10^ (95% CI 1.44 × 10^-10^ – 2.26 × 10^-10^), along with seven insertions and nine deletions (Table S5). Significantly reduced BPSs were found in protein-coding genes and nonsynonymous sites than expected, suggesting a possible role of purifying selection in preventing the fixation of strong deleterious mutations during the MA process ^38^ of *D. shibae* DFL-12. For *R. pomeroyi* DSS-3 (=DSM 15171^T^), the mutation rate was reported in our previous study ^20^, but its generation time was not corrected with its death rate. Here, we measured its death rate and updated its μ to be (1.38 ± 0.85) × 10^-10^ (95% CI 1.17 × 10^-10^ – 1.61 × 10^-10^). In summary, the genomic mutation data of CHUG, *S. pontiacus* and *R. pomeroyi* are unbiased, whereas that of *D. shibae* may be slightly biased owing to a significant deficit in nonsynonymous mutations than expected (SI Results).

Among the four Roseobacter species (Fig. 2A), CHUG shows a significantly higher μ than the other three species (Wilcoxon–Mann–Whitney test, *p* < 0.001 in all three comparisons). Among the remaining species, *S. pontiacus* shows a significantly higher mutation rate than *D. shibae* and *R. pomeroyi* (*p* < 0.001 in both comparisons). No significant difference was found between *D. shibae* and *R. pomeroyi*. Our results show a negative correlation between genome size and μ among the Roseobacter lineages (dashed gray line in Fig. 2B [*r*^2^ = 0.992, slope = −2.895, s.e.m. = 0.019, *p* = 0.004]) according to a generalized linear model (GLM) regression. This relationship is not caused by shared ancestry, as confirmed by phylogenetic generalized least square (PGLS) regression analysis (solid blue line in Fig. 2B [*r*^2^ = 0.991, slope = −2.894, s.e.m. = 0.159, *p* = 0.003]). This conclusion remains robust by leaving out *D. shibae*.

**Fig. 2.**
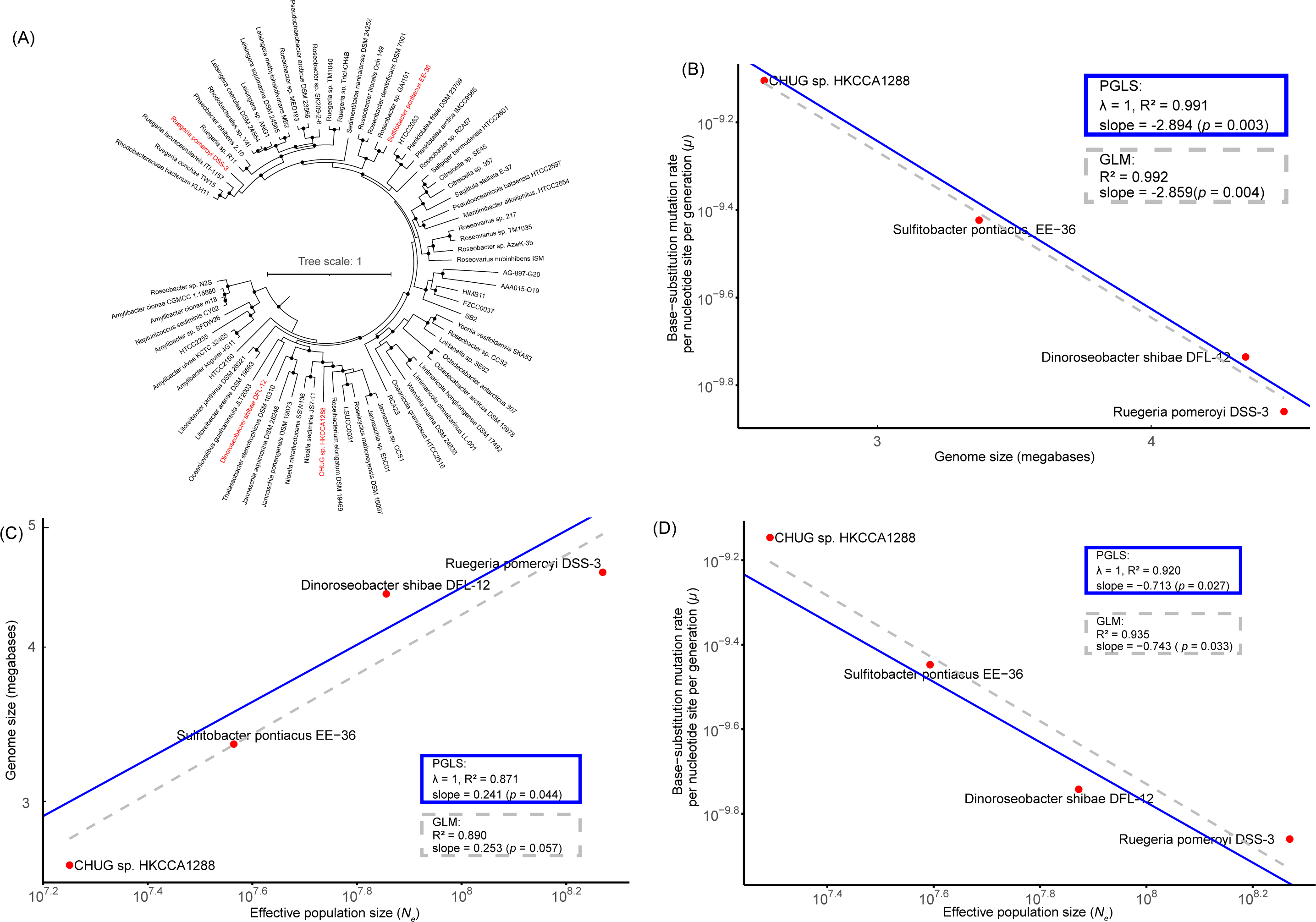
Phylogenomic tree of the Roseobacter group and scaling relationships between genome size, genomic mutation rate (base-substitution mutation rate per cell division per nucleotide site; μ) and effective population size (*N_e_*) for the studied Roseobacter species. All trait values were logarithmically transformed. **(A)** The maximum likelihood phylogenomic tree of representative Roseobacter genomes based on 120 conserved bacterial genes, with the four studied Roseobacter species marked in red. **(B-D)** Scaling relationships involving μ, *N_e_*, and genome size across four phylogenetically diverse lineages within the marine Roseobacter group, with all trait values logarithmically transformed. The dashed grey lines and solid blue lines represent the generalized linear model (GLM) and phylogenetic generalized least square (PGLS) regression, respectively.

### Genome sizes of the Roseobacter group members from surface oceans scale positively with effective population sizes

We calculated the *N_e_* of CHUG according to the aforementioned equation π_S_ = 2 X N_e_ X µ. Here, π_S_ estimation requires the delineation of the population boundary. In many previous studies involving bacterial *N_e_*estimates, π_S_ was estimated based on a nominal bacterial species ^39,40^. However, *N_e_* should be estimated based on neutral genetic diversity of a well-mixed population, whereas a nominal bacterial species is commonly comprised of structured populations ^9,41^. The recently available tool PopCOGenT defines a bacterial population whose members recombine more frequently than bacteria involving different populations ^42^, thereby rendering itself a powerful method for bacterial *N_e_* estimation. It was shown to outperform earlier methods and was used to delineate population boundaries of many prokaryotic species for *N_e_* estimation ^12^. We therefore used PopCOGenT to delineate populations for CHUG based on the published genomes of 39 isolates we sampled mostly from brown algae ambient seawater ^25,26^ as well as newly sequenced genomes of seven isolates from more diverse marine ecosystems such as regular coastal seawater and coral ambient seawater (Fig. S2). This approach led to the identification of two populations, one containing 21 non-redundant members (CHUG_MC0) with a genomic median π_S_ of 0.043 translated to *N_e_* of 2.74 X 10^7^, and the other containing four members (CHUG_MC1) with a median π_S_ of 0.013 and *N_e_* of 8.27 X 10^6^. This gave an average *N_e_* of (1.78 ±1.35) X 10^7^.

Using the same method, we estimated the mean *N_e_* of the two populations of *S. pontiacus* (Fig. S3 and Table S9) and median *N_e_*of five populations related to *R. pomeroyi* (Fig. S4 and Table S9) as (4.01 ± 3.73) X 10^7^ and 1.51 X 10^8^, respectively. There were only four *Dinoroseobacter* genomes publicly available, two (*D. shibae* DSM 112351 and *D. shibae* DFL-12 (= DSM 16493)) of which were derived from the same strain, and we extended the dataset by sequencing four additional strains. However, seven of them fall into a clonal complex (see Method) and are thus redundant. As a result, *N_e_* of this species was estimated by using only two non-redundant genomes (*D. shibae* PD6 and *D. shibae* DFL-12), which gave 7.35 X 10^7^ (Fig. S5) and remains to be updated in the future when more non-redundant strains become available.

The differences of *N_e_* across the three genome size categories (2-3, 3-4, 4-5 Mbp) of these surface ocean Roseobacter lineages are statistically significant. Specifically, CHUG has a significantly lower *N_e_*than the other three lineages (Wilcoxon–Mann–Whitney test, *p* < 0.001 in all cases), and *S. pontiacus* has a significantly lower *N_e_* than *Ruegeria* sp. and *D. shibae* (Wilcoxon–Mann–Whitney test, *p* < 0.001 in all cases). Our results show that genome size correlates positively with *N_e_*among these Roseobacter lineages (Fig. 2C). At first glance, this correlation is not significant according to the generalized linear model (GLM) regression analysis (dashed gray line in Fig. 2C [*r*^2^ = 0.890, slope = 0.253, s.e.m. = 0.063, *p* = 0.057].

However, GLM is not appropriate in this case because there is a strong phylogenetic effect on the relationship between these two traits (indicated by the λ value of 1 in blue box of Fig. 2C). By controlling for the phylogenetic signal, the phylogenetic generalized least square (PGLS) supports significantly positive relationship (solid blue line in Fig. 2C [*r*^2^ = 0.871, slope = 0.241, s.e.m. = 0.052, *p* = 0.044]).

### Increased mutation rate correlates with increased power of genetic drift among the Roseobacters

According to the GLM regression, we found a significantly negative relationship between μ and *N_e_* across the four Roseobacter lineages (dashed gray line in Fig. 2D [*r*^2^ = 0.935, slope = −0.743, s.e.m. = 0.139, *p* = 0.033]). Although the observed relationship was impacted by phylogenetic effect (indicated by λ value at 1), it remains robust after controlling for shared ancestry, as shown by PGLS regression analysis (solid blue line in Fig. 2D [*r*^2^ = 0.920, slope = −0.713, s.e.m. = 0.120, *p* = 0.027]).

### The scaling relationships between genome size, mutation rate, and effective population size are robust across prokaryotes

Based on 31 prokaryotic species (mostly bacteria, mostly from non-marine habitats) with µ determined with the MA/WGS strategy (Table S10) and *N_e_* calculated with the same approach as presented here, Chen et al. (2022) reported that both μ and the genome-wide mutation rate (*U_P_*, a proxy for deleterious mutation load of a genome ^11^) scales negatively with *N_e_*. They also reported a negative scaling relationship between genome size and μ, but they found the negative correlation between genome size and *N_e_* was not significant by both GLM and PGLS regression analyses ^12^. By including the new data of the three Roseobacter lineages determined here, we confirmed the first three correlations (Fig. S6A&B&C). Intriguingly, we found a significant positive scaling relationship between genome size and *N_e_*after controlling for the common ancestry (PGLS, solid blue line in Fig. S6D [*r*^2^ = 0.251, slope = 0.147, s.e.m. = 0.048, *p* = 0.005]).

## Discussion

The accepted genome streamlining theory attributes the process of genome reduction to selection for efficient use of limited nutrients ^5^. For this reductive process to be visible to natural selection, the lineages undergoing genome streamlining must have greater *N_e_* than their co-occurring sister lineages that carry larger genomes ^6^. To study streamlining process in surface ocean bacterioplankton cells, *N_e_* is the key parameter and bacterioplankton lineages found in surface oceans should be compared. The marine Roseobacter group provides a unique opportunity to test it. On one hand, group members that co-exist in surface oceans span a wide range of genome sizes. On the other hand, many Roseobacter populations are primarily associated with nutrient-rich marine habitats such as coral holobionts ^43^ and benthic environments ^44^. These species are not appropriate targets to study the streamlining process because their growths are less limited by nutrients and their genome sizes are additionally shaped by characteristics of their habitats (e.g., host immune responses for host-associated Roseobacters) that are very different from those of surface oceans. In the case of the four Roseobacter lineages studied here, CHUG is featured by decoupling itself from marine eukaryotic phytoplankton groups ^26^, whereas *Dinoroseobacter shibae* is found to be primarily associated with marine phytoplankton ^45^. Commonly found in between the two oceanic niches is *Ruegeria pomeroyi* ^46^.

Less is known for *Sulfitobacter pontiacus*, though other *Sulfitobacter* species are commonly associated with phytoplankton ^47,48^. Although the difference in their ecological strategies (phytoplankton-associated versus free-living) are relevant to the difference in their genome sizes, these lineages remain valuable for testing the streamlining theory because they are all commonly found in surface ocean habitats and are generally subjected to nutrient limitation. Simulations with an agent-based model showed that carbon limitation at geological timescales is the primary force that shapes the high genomic G+C content of *R. pomeroyi* DSS-3 ^49^.

A surprising result from the present study is that the *N_e_*of the surface ocean Roseobacter lineages scales positively with their genome sizes (Fig. 2C). This is unexpected because it reverses the trend predicted by the streamlining selection theory. As the strength of genetic drift is the inverse of *N_e_*, this finding means that genetic drift becomes increasingly powerful in bacterioplankton species that carry increasingly small genomes. For selection to eliminate a deleterious mutation and promote a beneficial mutation, the mutation needs to be sufficiently deleterious or beneficial (i.e., the absolute value of selection coefficient *s*, |*s*|, is large enough) to overcome the power of genetic drift (1/*N_e_*), or the condition |*s*| > 1/*N_e_* needs to be fulfilled. For a population with a decreased *N_e_*, more deleterious mutants are expected to be fixed and more beneficial variants will be lost by chance, rendering natural selection less effective. Therefore, our finding suggests that natural selection becomes less effective for the surface ocean Roseobacter populations with decreasing genome sizes.

For genetic drift to play a role in bacterial genome reduction, it needs to work with certain mutational processes. Most mutations are deleterious and purged primarily by purifying selection and recombination ^50,51^. In a small population with limited opportunities for recombination, deleterious mutations are accumulated by random and irreversible loss of genotypes that are depleted with deleterious mutations and by random fixation (i.e., 100% in frequency) of genotypes that are loaded with abundant deleterious mutations. This process is known as “Muller’s ratchet” ^52^. A direct effect of Muller’s ratchet in bacteria is genome reduction due to accumulation of irreversible disabling mutations that lead to pseudogenization and gene loss. Because obligate endosymbiotic bacteria are featured by very small population sizes with extremely low recombination rates, their reductive evolution is generally driven by Muller’s ratchet ^53,54^.

As marine bacterioplankton lineages have much higher recombination rates than obligate endosymbionts, their evolution was thought to be unlikely to be driven by Muller’s ratchet ^55^. Although this is true in the overall well-connected oceans, recombination may be restricted in a frozen ocean of the globe in some unusual geological periods of the Earth. A prominent example is that in the global icehouse climate conditions during the Neoproterozoic Snowball Earth that comprises Sturtian (approx. 717–659 Ma) and Marinoan (approx. 645–635 Ma) glaciations, *Prochlorococcus* cells were likely restricted to a few biotic refugia such as cryoconite holes and sea-ice brine channels ^56^. Such highly patchy and disconnected habitats likely created little opportunity for recombination, which was evidenced by extremely few gene acquisitions at that stage ^57^. Additionally, *Prochlorococcus* at that time experienced severe population bottlenecks, bringing its *N_e_* down to 10^4^-10^5^ or even lower according to simulations by an agent-based model ^57^. The very small *N_e_*and the very rare recombination support an escalated role of Muller’s ratchet as a driver of the *Prochlorococcus* evolution at that time. As the *Prochlorococcus* major genome reduction event, where ∼30% of the genomic DNA was eliminated, also occurred at that time, it is natural to come up with a new theory that Muller’s ratchet likely acted as a main mechanism of the historical genome reduction in *Prochlorococcus* ^57^.

Unlike *Prochlorococcus* that are found exclusively between 40° N and 40° S and thus vulnerable to global icehouse climate, oceanic members of the Roseobacter group are globally distributed and generally enriched in ocean regions at middle and high latitudes ^14,15^, thereby more resistant to glaciation events and less likely to undergo severe population bottlenecks during historical glaciation events. For this reason, genome reduction in the oceanic members of the Roseobacter group is less likely to be driven by Muller’s ratchet.

Alternatively, genetic drift may lead to genome reduction through mutation rate increases. This theory further builds on two population genetic processes: increased strength of genetic drift leads to increased mutation rate, and mutation rate increase leads to genome reduction. If mutation rate exceeds selection coefficient of a gene, purifying selection may not be able to keep the gene ^58,59^. With increasing mutation rates, more non-essential genes are lost and genome reduction occurs. This explains why mutation rate increase leads to genome reduction. In fact, a potential role of mutation rate increases in marine bacterial genome reduction has been repeatedly hypothesized for *Prochlorococcus* and SAR11 ^8,59–61^. The present study is the first that experimentally validated this long-lasting hypothesis, though genome-reduced Roseobacters, rather than *Prochlorococcus* and SAR11, were used in the analysis. A theoretical model predicts that bacterial genome size is reduced by 30% in response to mutation rate increase by 10 times ^59^. It seems that our empirical data from natural Roseobacter lineages is broadly consistent with this prediction.

However, the population genetic process giving rise to increased mutation rate in bacteria has been more disputable. On one hand, the “mutator theory” ^62–64^ favors natural selection as the primary force to boost mutation rate; it posits that a high mutation rate may provide transient advantages for prokaryotes in a changing environment as it increases the opportunity to gain beneficial mutations. On the other hand, the “drift-barrier” model posits that natural selection favors low mutation rates and acts to increase replication fidelity to a threshold beyond which genetic drift can overcome the effects of natural selection and thus fitness advantages start to decrease. The drift-barrier model favors a primary role of genetic drift in increasing mutation rates and predicts an inverse relationship between *N_e_* and μ ^11,40^. It gained a great success in explaining why eukaryotes (generally having small *N_e_*) have higher mutation rates than prokaryotes (generally having large *N_e_*) ^11,40^. From within bacteria, the model was originally skeptical when genome-reduced marine bacterioplankton lineages were concerned. These marine bacteria, especially *Prochlorococcus* and SAR11, were believed to have unusually large *N_e_*^6,65^ and assumed to have high μ (because of losses of important repair systems) ^8,59–61^. Not until recently when the *N_e_* and μ of a genome-reduced *Prochlorococcus* strain became available, the drift-barrier model received stronger support from within bacteria ^12^. The inclusion of the three Roseobacter species determined here, especially the CHUG, continues to support the negative scaling relationship between *N_e_* and µ (Fig. S6C), thus strengthening the drift-barrier theory. A gross relationship across deeply-branching bacterial lineages means that genetic drift is a universal rule dictating the evolution of mutation rates in bacteria. However, it does not mean that this rule is necessarily manifested by an analysis of bacterial species from a certain habitat because the pattern could be shadowed by selective pressure imposed by the particular environment. It is therefore remarkable to observe the inverse scaling relationship between *N_e_* and µ holds when only the four Roseobacter species were compared (Fig. 2D), supporting the idea that genetic drift drives mutation rate increases in the studied Roseobacters.

Our study shows that genetic drift and mutation rate increases are the two primary population genetic mechanisms that mediate genome reduction in surface ocean Roseobacter lineages. We also show that mutation rate increase itself is driven by genetic drift. Taken together, we are able to conclude that genetic drift is the ultimate mechanism for genome reduction in the studied marine bacteria. Our new result contributes to the ongoing discussion on the population genetic mechanisms giving rise to the small genomes in marine bacterial cells that dominate the marine bacterioplankton communities. It provides an alternative view to the prevailing streamlining selection theory, but the presented evidence from the studied surface ocean Roseobacter lineages, several of which are commonly associated with phytoplankton where nutrients are more available than in the bulk seawater, is not sufficient to reject the streamlining selection theory. The genome streamlining process is deemed to be more efficient in bacterioplankton cells that are found in highly oligotrophic and stratified oceans such as the ocean gyres where nutrients are extremely depleted, and thus the streamlining selection theory is believed to work best in explaining the genome reduction process in those bacterioplankton lineages. Future work should be focused on comparing lineages that dominate the bacterioplankton communities in oligotrophic oceans.

## Data and code availability

All the datasets generated, analyzed and presented in the current study are available in the Supplementary Information. Raw reads of the surviving lines of three Roseobacter lineages are available at the NCBI SRA under accession no. PRJNA1067981. The closed genome of *S. pontiacus* EE-36 are available at the NCBI GenBank under accession no. SAMN39584998. All the scripts are deposited at https://github.com/Xiaojun928/Genome_reduction_bacterioplankton.

## Author contribution

H.L. conceived and designed the study. X.W. performed the bioinformatics. H.L. and X.W. wrote the paper. M.X. and K.H. performed the CHUG MA experiment, M.X. measured the death rates of all MA experiments, validated the mutations of CHUG, and constructed DNA library for genome sequencing. Y.S. and X.C. performed the *S. pontiacus* EE-36 MA experiments. Y.S. assembled the complete genome of *S. pontiacus* EE-36. S.Z. and H.W. conducted the *D. shibae* DFL-12 MA experiments. X.C. and M.X. cultured new CHUG isolates. V.R., T.B., J.P. and I.W.D contributed the four new *D. shibae* strains. Z.S. and X-H.Z. contributed five and five *S. pontiacus* strains, respectively. M.X, Y.S. T.B., J.P. and I.W.D edited the manuscript. I.W.D provided substantial comments that significantly improved this manuscript.

## Supporting information

Supplemental mathods and results

## Acknowledgements

We thank Danli Luo for the isolation of two CHUG members from ambient water of corals, Ho Ching Leung for helping prepare the CHUG MA experiments, and Yang Qian for his assistance in the isolation of some CHUG strains. This work is supported by the Hong Kong Research Grants Council General Research Fund (14110820) and the Hong Kong Research Grants Council Area of Excellence Scheme (AoE/M-403/16). X.W. was supported by the China Postdoctoral Science Foundation (2022M722220). T.B, J.P and I.W.D were supported by the Deutsche Forschungsgemeinschaft within the Collaborative Research Centre TRR51 Roseobacter.

