## Supplemental Figures 1-6 for "A neutral process of genome reduction in marine bacterioplankton"

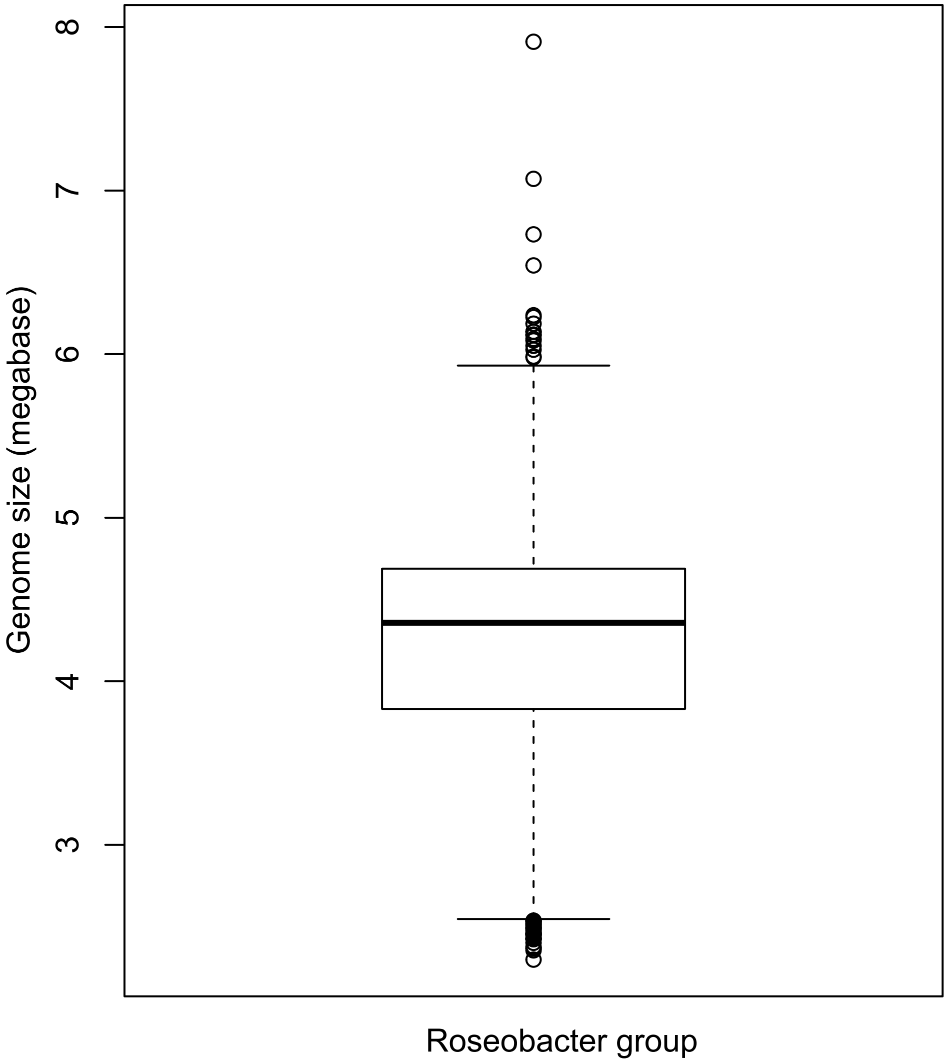


Fig. S1. Box plot for genome size variation of public Roseobacter strains. The center line denotes the median value (50th percentile), and the box spans from the 25th to the 75th percentiles of the dataset. The black whiskers mark the 5th and 95th percentiles, and values beyond these upper and lower bounds are outliers marked with black circles.

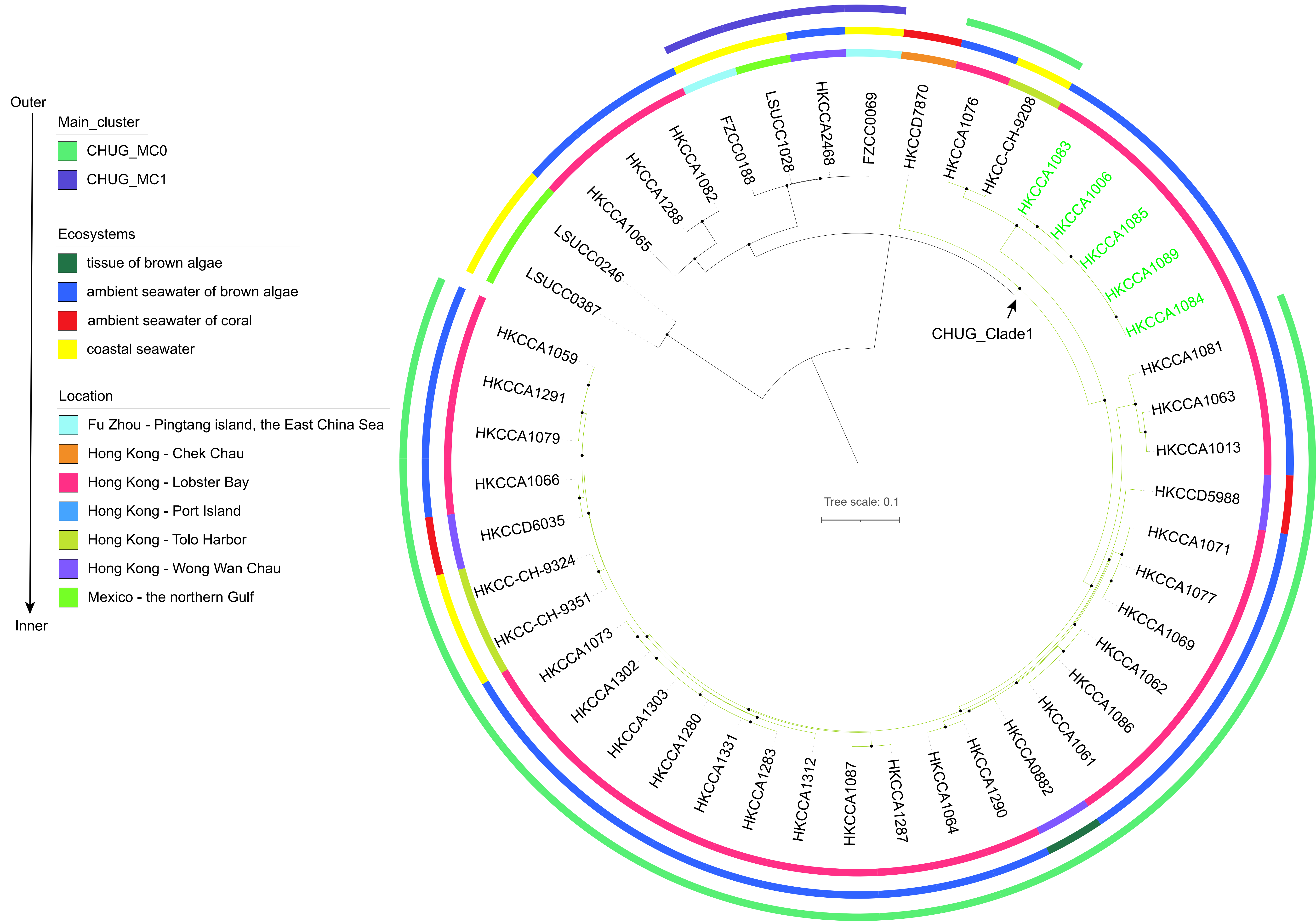


Fig. S2. The maximum-likelihood phylogenomic tree of 46 CHUG genomes and sampling information mapped to the tree. Solid circles at the internal nodes indicate the frequency of the group defined by that node is at least 90% in the 1,000 bootstrapped replicates. The tree is visualized and annotated with iTOL. Moving from the outer to inner rings: (1) PopCOGenT delineated populations with at least three non-redundant members; (2)-(3) the sampling niche and location from which each CHUG strain was isolated. The branches highlighted in green represent CHUG_Clade1, which consists of CHUG_MC0 and a clonal complex of five genomes (in green).

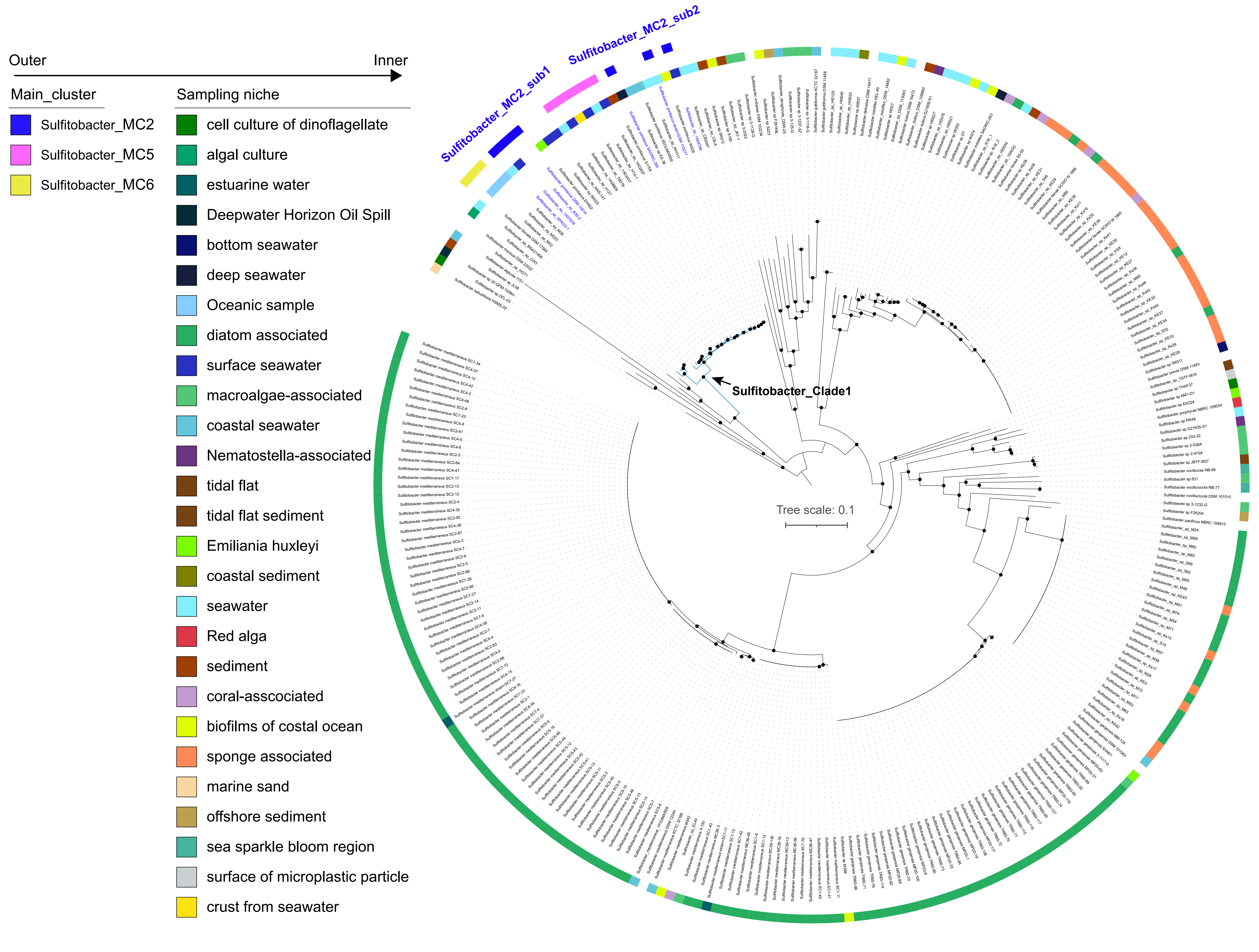


Fig. S3. The maximum-likelihood phylogenomic tree of 279 *Sulfitobacter* genomes and sampling information mapped to the tree. Solid circles at the internal nodes indicate the frequency of the group defined by that node is at least 90% in the 1,000 bootstrapped replicates. The tree is visualized and annotated with iTOL. Moving from the outer to inner rings: (1) PopCOGenT delineated populations with at least three non-redundant members and the majority originating from surface seawater; (2) sampling niche where each *Sulfitobacter* strain was isolated. The branches in blue correspond to *Sulfitobacter*_Clade1, which consists of *Sulfitobacter*_MC2 and additional 24 descendants. The seven strains constituting *Sulfitobacter*_MC2 are marked in blue, which were not used for $\pi_s$ and $N_e$ estimation because they did not form a monophyletic group.

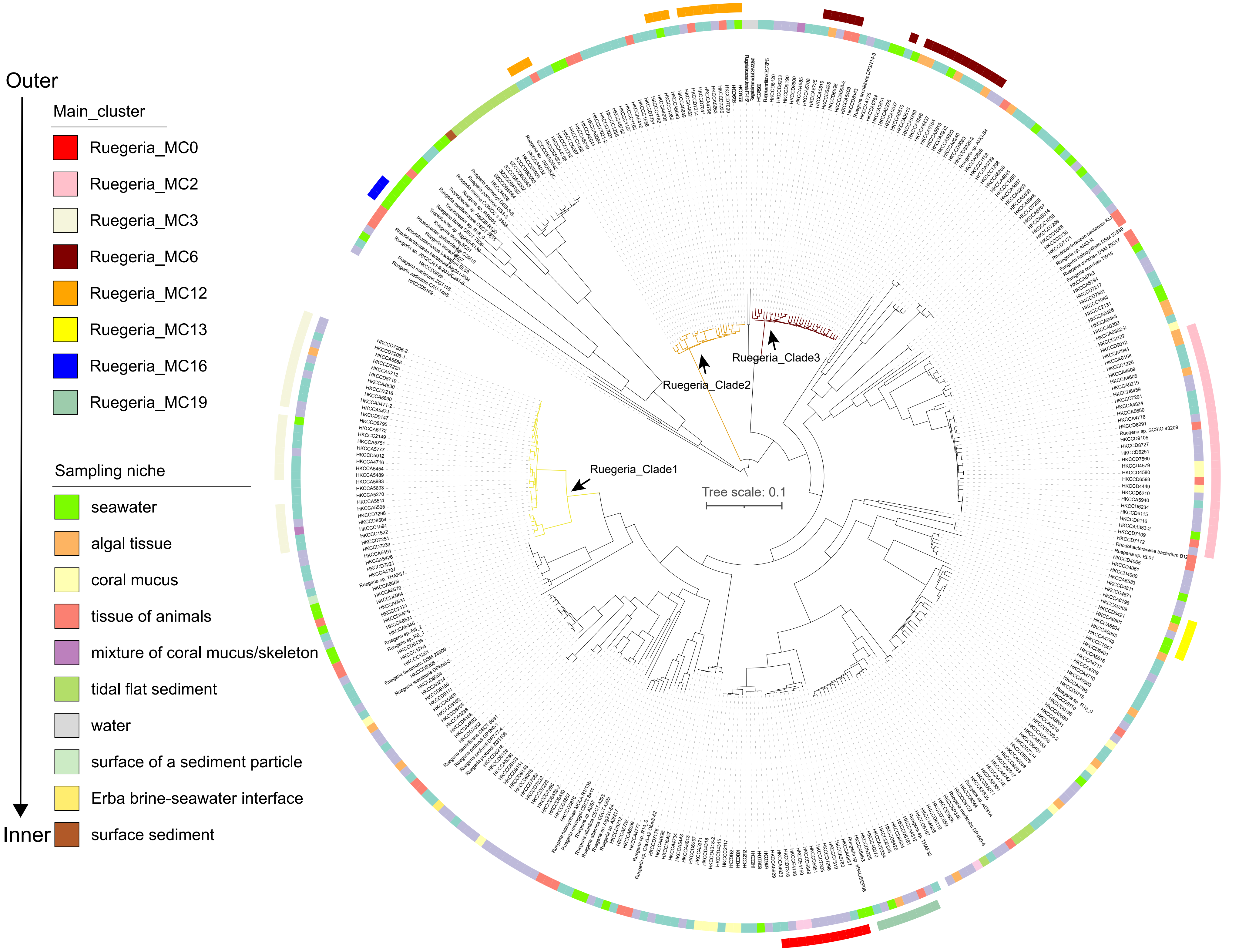


Fig. S4. The maximum-likelihood phylogenomic tree of 353 *Ruegeria* genomes and sampling information mapped to the tree. Solid circles at the internal nodes indicate the frequency of the group defined by that node is at least 90% in the 1,000 bootstrapped replicates. The tree is visualized and annotated with iTOL. Moving from the outer to inner rings: (1) PopCOGenT delineated populations with at least three non-redundant members and at least one strain derived from surface seawater; (2) sampling niche where each *Ruegeria* strain was isolated. The branches in yellow, orange, and brown are *Ruegeria*_Clade1, *Ruegeria*_Clade2, and *Ruegeria*_Clade3 respectively, corresponding to the extended clade of *Ruegeria*_MC3, *Ruegeria*_MC12, and *Ruegeria*_MC6 respectively. These three populations were not used for $\pi_S$ and $N_e$ estimation as they each did not form a monophyletic group.

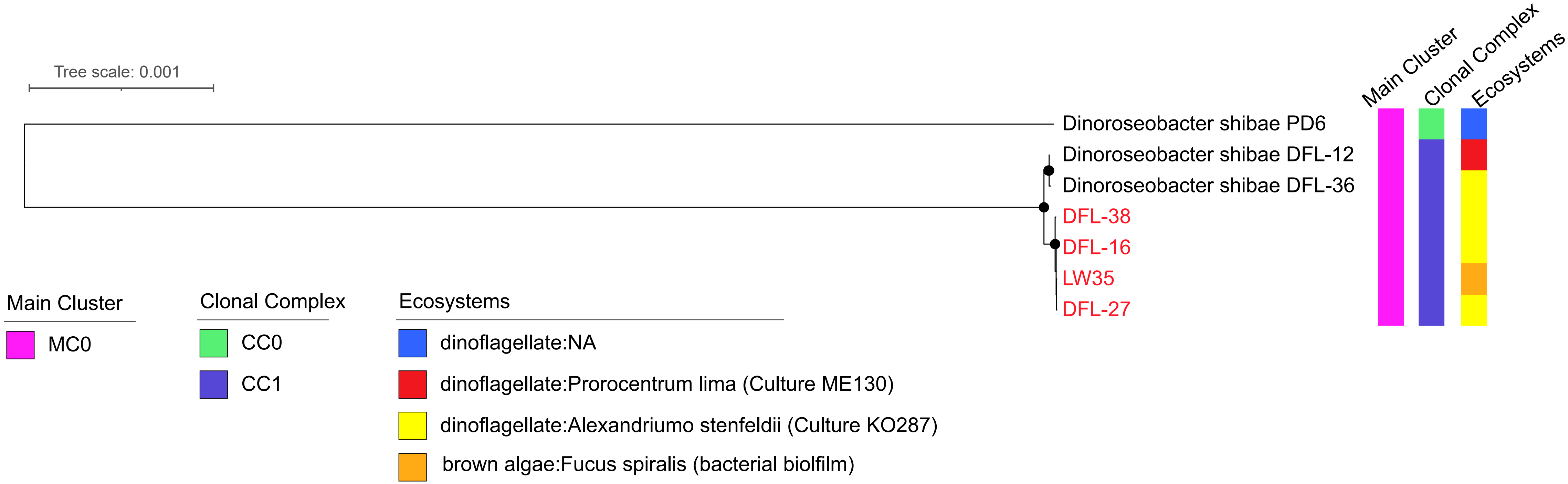


Fig. S5. The maximum-likelihood phylogenomic tree of the seven *Dinoroseobacter* genomes and sampling information mapped to the tree. Solid circles at the internal nodes indicate the frequency of the group defined by that node is at least 90% in the 1,000 bootstrapped replicates. The tree is visualized and annotated with iTOL. Strains isolated in this study are marked in red. Moving from the outer to inner rings: (1) PopCOGenT delineated population, (2) PopCOGenT delineated clonal complex, (3) the sampling niche where each *Dinoroseobacter* genome was isolated.

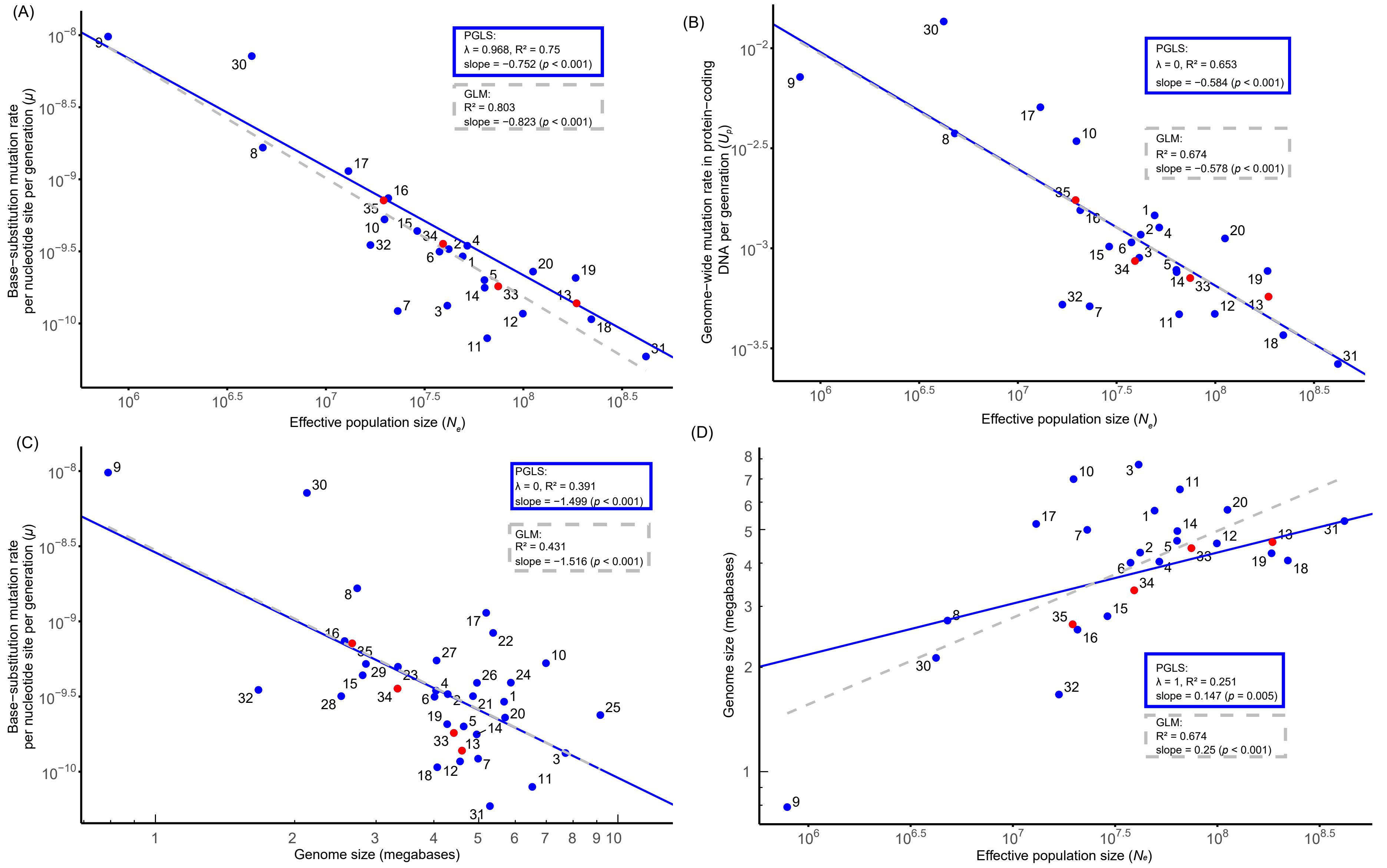


1 Agrobacterium tumefaciens C58
2 Bacillus subtilis NCIB 3610
3 Burkholderia cenocepacia HI2424
4 Caulobacter crescentus NA1000
5 Escherichia coli K 12 MG1655
6 Haloferax volcanii DS2
7 Janthinobacterium lividum KBS0711
8 Lactococcus lactis DSMZ20481
9 Mesoplasma florum L1
10 Mycobacterium smegmatis MC2 155
11 Pseudomonas aeruginosa PA14
12 Rhodobacter sphaeroides ATCC17025
13 Ruegeria pomeroyi DSS 3
14 Salmonella enterica
15 Staphylococcus aureus ATCC 25923
16 Staphylococcus epidermidis ATCC 12228
17 Teredinibacter turnerae
18 Vibrio cholerae 2740 80
19 Vibrio fischeri ES114
20 Vibrio shilonii AK1
21 Arthrobacter sp. KBS0703
22 Colwellia psychrerythraea 34H
23 Deinococcus radiodurans R1
24 Flavobacterium sp. KBS0721
25 Gemmata obscuriglobus DSM5831
26 Kineococcus radiotolerans SRS30216
27 Leeuwenhoekiella sp. ZYFB001
28 Micrococcus sp. KBS0714
29 Nonlabens sp. SY33080
30 Thermococcus eurythermalis A501
31 Photorhabdus luminescens ATCC29999
32 Prochlorococcus marinus AS9601
33 Dinoroseobacter shibae DFL-12
34 Sulfitobacter pontiacus EE-36
35 CHUG sp. HKCCA1288

Fig. S6. Scaling relationships involving the base-substitution mutation rate per cell division per nucleotide site ($\mu$), and genome-wide mutation rate per cell division per genome ($U_P$), estimated effective population size ($N_e$), and genome size across 33 bacterial and two archaeal species. All trait values were logarithmically transformed. The mutation rates of species CHUG HKCCA1288, *S. pontiacus* EE-36, and *D. shibae* DFL-12 are from the present study, whilst that of the remaining species are collected from literature. All species in the Roseobacter group are denoted with red dots. Mutation rates of all 35 species were determined using the MA/WGS strategy. (A) $\mu$ scales negatively with Ne. (B) $U_P$ scales negatively with $N_e$. (C) Genome size scales negatively with $\mu$. (D) Genome size scales positively with $N_e$. Numbered data points 21–29 are not shown in A, B and D owing to the lack of population-level datasets required for the estimation of $N_e$. The dashed grey lines and solid blue lines represent the GLM and PGLS regression, respectively.
