## Supplemental mathods and results for "A neutral process of genome reduction in marine bacterioplankton"

**This file includes the following items:**

Supplementary Materials and Methods

Supplementary Results

Supplementary References

### Supplementary Materials and Methods

#### Mutation accumulation experiment and genome sequencing of mutant lines

We performed mutation accumulation (MA) experiments on four strains in the Roseobacter group. They are a representative CHUG strain HKCCA1288 isolated from the ambient seawater of *Sargassum* *hemiphyllum* in Hong Kong ^1^, *Sulfitobacter* *pontiacus* EE-36 from salt marsh at the coast of Georgia, USA ^2^, *Dinoroseobacter shibae* DFL-12 from the dinoflagellate *Prorocentrum lima* ^3^, and *Ruegeria pomeroyi* DSS-3 from coastal water of Georgia, USA ^4^. Except for DSS-3 published in an earlier study ^5^, all were done here. For the CHUG lineage, 200 independent MA lines were initiated from a single founder colony of strain HKCCA1288. Throughout the experiment, a single colony from each MA line was transferred to a fresh 2216E marine agar (BD Difco, USA) plate and incubated at 28℃ every four days. This repeated single-colony bottlenecking process ensures that mutations accumulate in an effectively neutral fashion. After each transfer, the original plate was retained as a backup plate at 4˚C. If the new plate got contaminated or a single colony could not be picked, a single colony was picked from the most recent backup plate kept at 4˚C. Such situations arose less than 20 times during the entire experiment. The MA propagation was completed following 64 transfers in 252 days, and eight MA lines were lost during the MA process. Genomic DNA of the 192 survived MA lines was extracted using TIANamp Genomic DNA Kit (DP304, Tiangen, Beijing, China). DNBSEQ 150-bp paired-end sequencing (BGI, China) was applied to each of the 192 MA lines, with an average depth of sequencing coverage being 199 ± 174.

For the *Sulfitobacter* lineage, 111 independent MA lines were initiated from a single founder colony of strain *S. pontiacus* EE-36. The propagation procedure is similar to that of strain HKCCA1288 except that transfer was performed daily, and the MA propagation was completed following 188 transfers in 188 days. In total, 110 MA lines survived during the MA process and genomic DNA of each survived line was extracted using TIANamp Genomic DNA Kit (DP304, Tiangen, Beijing, China). The 251-bp paired-end Illumina (Illumina HiSeq platform) sequencing was applied to each of the 110 MA lines, with an average depth of sequencing coverage being 130 ± 50. In our first sequencing, raw reads in most samples are in poor quality. In detail, 29 MA lines showed low number of reads, or low coverage depth or breadth after quality control. Despite the remaining 81 MA lines showed over 95% coverage breadth, they were influenced by biased sequencing errors, e.g., nucleotides G, C, T have more than six times error rate than nucleotide A. Thus, we randomly selected 58 out of the 81 lines for resequencing and used them for mutation calling owing to high sequencing quality.

For the *Dinoroseobacter* lineage, 220 MA lines were initiated from a single founder colony of strain *D. shibae* DFL-12. The MA propagation procedure is similar to that of strain HKCCA1288 except that transfer was performed every three days, the propagation was completed following 75 transfers in 222 days, and cells were incubated in dark. In total, 187 MA lines survived during the MA process and genomic DNA of each survived line was extracted using TIANamp Genomic DNA Kit (DP304, Tiangen, Beijing, China). DNBSEQ 150-bp paired-end sequencing (BGI, China) was applied to each of the 187 MA lines, with an average depth of sequencing coverage being 136 ± 53.

#### Estimation of generation time, death rate, total cell divisions, and effective population size of MA lines

To estimate the total cell divisions during the MA experiment, the number of cell divisions per transfer (*D*) was measured for the three strains (HKCCA1288, *S. pontiacus* EE-36, and *D. shibae* DFL-12) respectively using the live cell density (*d*). Briefly, an entire single colony was cut from each of the ten randomly selected MA lines and then transferred to 1 mL autoclaved seawater. To measure *d*, these samples were vortexed, serially diluted, spread on 2216E agar plates, and cultured at 28 °C until counting. The *d* value of each sample was then calculated from the viable cell counts multiplied by the dilution factor. However, not all cells survive in the plate, and the dead cells would not contribute to the *d* value, which, if not taken care of, would lead to underestimation of *D*. Thus, we corrected *D* value using the ratio of live cells to total cells (*r*) according to the formula $D={log}_{2}(d/r)$ ^6^. To measure *r*, 150 μL of each sample was separately stained by 1,000 times diluted SYBR Green I (10 mg/mL, Invitrogen, USA) for both live and dead cells and by Prodium Iodide (20mM, Invitrogen, USA) for dead cell only. The *r* value was then determined with flow cytometry (BD FACSVerse, BD Biosciences, USA). The *D* value of *R.* *pomeroyi* DSS-3 in previous publication ^5^ has not been corrected using death rate, we thus used the same assay to measure its death rate and calculated its *D* value. The total number of generations for each MA line was calculated by the average number of cell divisions per transfer (Table S11) multiplied by the total number of transfers.

The effective population size (*N_e_*) of an MA line was calculated based on the harmonic mean ^7–10^ according to the following equation:

$$N_{e}= \frac{n+1}{\sum_{i=0}^{n} \frac{1}{2^{i}}}$$

, where $n$is the number of cell divisions from the initial population (with a population size of 1) to the final population represented by a single colony of the target strain and measured by flow cytometry. Here, $n$ is 23.20 ± 0.74, 21.82 ± 0.58, 19.29 ± 1.28, and 22.80 ± 1.31 according to the logarithm base 2 of the final population size of HKCCA1288, *S. pontiacus* EE-36, *R. pomeroyi* DSS-3, and *D. shibae* DFL-12, respectively. Accordingly, *N_e_* was calculated to be 12, 11.5, 10, and 12, respectively, suggesting a negligible role of natural selection during MA processes.

#### Reference genomes

To identify mutations in each MA experiments, the closed reference genomes of HKCCA1288 and *D. shibae* DFL-12 were collected from NCBI GenBank. Strain *D. shibae* DFL-12 carries 47 pairs of repetitive genomic regions (nucleotide identity >99.9%) each over 150 bp, which may interfere with mutation calling, so two reference genomes of *D. shibae* DFL-12 were used in mutation calling. One is the original genome, and another is the genome with repetitive genomic regions masked by replacing nucleotides with ‘N’ bases. In terms of *S. pontiacus* EE-36, a closed genome was not publicly available, it was sequenced with PacBio Sequel platform and assembled with Unicycler v0.4.6 ^11^ to obtain a complete and closed genome. Protein-coding genes of HKCCA1288 and *D. shibae* DFL-12 were downloaded from the NCBI GenBank, and those of *S. pontiacus* EE-36 genome were predicted with Prokka v1.12 ^12^.

#### Mutation calling and mutation rate estimation

For all three MA experiments, mutations were identified as described previously ^13^. In brief, raw reads were processed by Trimmomatic v0.32 ^14^ to remove adaptors and trim low-quality bases with the parameters of ‘ILLUMINACLIP: 2:30:10:1:true SLIDINGWINDOW:4:15 MAXINFO:40:0.9 MINLEN:40’. Then the paired-end reads were individually mapped to the corresponding reference genome using BWA-mem v.0.7.17 ^15^. The output was parsed and converted to SAM format with SAMTOOLS v.1.4.1 ^16^. Next, these SAM files were processed using Picard MarkDuplicates (http://broadinstitute.github.io/picard/) to remove duplicate reads which may arise during the sequencing process, like polymerase chain reaction (PCR) duplication artefacts.

BaseRecalibrator in GATK-4.0 ^17^ was used for base quality recalibration to adjust the base quality score affected by systematic technical errors. Then base substitutions and small indels were called using HaplotypeCaller implemented in GATK-4.0 ^17^. Variants were further filtered with standard parameters recommended by GATK best practice, except that the Phred-scaled quality score QUAL > 100 and root-mean-square mapping quality MQ > 59 were set by following previous studies ^18–20^. A minimum of 10 reads and 80% of reads in a line were required to call the line-specific consensus nucleotide at a site. An in-house script was implemented to identify genes showing significant enrichment of mutations (bootstrap test, *p* < 0.05 for each gene).

To confirm the called mutations, three base-substitution mutations and four indels were randomly sampled from six CHUG MA lines and validated using PCR (Table S2). In brief, primers were designed to amplify a region of DNA covering the mutation site, approximately 600 bp in length including 300 bp each at the upstream and downstream to the mutation site, and then Sanger sequencing was performed on the PCR products (Table S12).

For all three MA experiments, base-substitution mutation rate was calculated using the following method. First, the base-substitution mutation rate per nucleotide site per cell division (μ_bs_) for each line was calculated according to the equation $\mu= \frac{m}{ng}$, where $m$ is the number of observed base substitutions, $n$ is the number of nucleotide sites analyzed, and $g$ is the mean number of cell divisions estimated covering the entire MA process. The standard error of $\mu$ across all MA lines was calculated as ${s.e.}_{pooled}= \frac{s}{\sqrt{N}}$, where $s$ is the standard deviation of the mutation rate across all lines and *N* is the number of lines analyzed.

For *D. shibae* DFL-12, two additional procedures were followed to estimate its mutation rate. First, some MA lines of *D. shibae* DFL-12 showed high enrichment of reads in some genomic regions but low read coverage in others, so both coverage depth and breadth were estimated for individual MA lines based on the SAM files using SAMTOOLS v1.4.1 ^16^. Consequently, we removed 38 MA lines. Of these, 36 lines each had only ~20% of their genomic regions covered with raw reads. In the other two MA lines, although ~80% of the genomic regions were covered, obvious systematic sequencing errors were found in their raw reads according to GATK Base Quality Score Recalibration. Hence, we kept the remaining 149 MA lines for further analysis, as their reads covered more than 95% of the genomic regions with coverage depth over 10× and show no systematic sequencing errors. Second, two reference genomes (i.e., the original one and the one with repetitive regions masked) were used for mutation calling, yielding nearly the same mutation rates (Table S5 vs Table S7) and mutation spectrum (Table S6 vs Table S8). To exclude any effect of repetitive genomic regions, the results based on masked genome were presented in the main paper (Table S5 & Table S6).

#### Mutation rate correction

Generally, mutations occur randomly along the genomic regions, but 17, three and eight protein-coding genes showed an excess of mutations (bootstrap test, *p* < 0.05 for each gene) in the MA lines of HKCCA1288, *S. pontiacus* EE-36, and *D. shibae* DFL-12, respectively. In detail, the 17 genes of HKCCA1288 accumulated 43 base-pair substitutions (BPSs) and 72 insertions and three deletions, the three genes of *S. pontiacus* EE-36 harbored four BPSs and seven indels, and the eight genes of *D. shibae* DFL-12 had 78 BPSs and 64 indels. Besides, two ribosomal genes (16S rRNA and 23S rRNA genes) of HKCCA1288 accumulated 40 BPSs and two insertions. The clustering of mutations indicates either mutation hotspots or that mutations in these genes confer selective advantages under the experimental condition. To exclude the potential effect of positive selection and possible epistatic interactions between these mutations and mutations in other genomic regions, the mutation rate for each strain was re-estimated using the MA lines without any mutation occurring in those genes. The strains HKCCA1288, *S. pontiacus* EE-36, and *D. shibae* DFL-12 had 97, 51, and 62 remaining lines, respectively. They accumulated 270, 243, and 80 BPSs, respectively, which were used to re-calculate the mutation rates of the three strains. To test if using a different number of lines may affect mutation rate estimation, we subsampled 51 MA lines (according to *S. pontiacus* EE-36) from HKCCA1288 and *D. shibae* DFL-12 species, respectively. This process was repeated 100 times for each species. The mutation rate derived from each subsample of MA lines does not significantly differ from that obtained using all the MA lines, supporting the idea that mutation rate estimates are robust and not affected by using different numbers of MA lines across different species.

#### New and publicly available genome sequences of natural isolates of CHUG, Sulfitobacter, Ruegeria and Dinoroseobacter

Genomes of natural isolates were used to delineate population boundaries (see next section). For CHUG, we collected 39 publicly available genomes from NCBI GenBank database (on 14 July 2023) and sequenced the genomes of seven new strains isolated from Hong Kong coastal waters (see Table S9 for detailed sampling information). Among the new strains, two were isolated from coral ambient seawater with a modiﬁed marine basal medium recipe ^21^ and cultivated with 2216 marine agar (BD Difco, USA), and two were from brown algal ecosystem using the method published Chu et al., (2022). The remaining three CHUG strains were isolated from coastal seawater using the high-throughput cultivation (HTC) approach. In detail, 100 mL coastal seawater was sampled with the sterile flask and transported to the laboratory in the cool box. The concentration of bacterial cells in the seawater sample was then measured by flow cytometry. The culture medium contained 500 mL of autoclaved seawater, 500 μL of vitamin mixture and carbon source (including taurine 40 mM, D-glucose 10 g/L, D-ribose 10 g/L, pyruvate 10 g/L, N-acetylglucosamine 10 g/L, methionine 10 g/L, glycine 10 g/L, and ethanol 10 mL/L), and 50 μL of 100 mM K_2_HPO_4_ and 1M NH_4_Cl based on the ^23^. 2 mL of culture medium was distributed in each well of the 24-well plate. A certain volume of seawater sample was then added to each well to reach an average of four cells per well. The plates were cultured at 20˚C. The concentration of each well was measured monthly and a concentration of 10^5^ indicated “positive growth”. The biomass of the “positive growth” wells were collected by centrifuging at 10,000 rpm for one hour at 4˚C. The supernatant was then discarded and 20-50 μL TE buffer was added to resuspend the bacteria cells. This solution was boiled at 100˚C for 15 min to release DNA. The 16S rRNA gene was amplified using PCR with the boiled sample as the template and primers 27F (5'-AGAGTTTGATCCTGGCTCAG-3') and 1492R (5'-GGTTACCTTGTTACGACTT-3'). The CHUG members were determined if they were clustered with previous CHUG members in the phylogenetic tree based on 16S rRNA gene. For newly identified CHUG members, the bacteria cells were propagated in the 100 mL flask with fresh culture medium and the biomass was collected as above. The bacterial DNA was extracted by TIANamp Genomic DNA Kit (DP304, Tiangen, Beijing, China) and used for whole genome sequencing with DNBSEQ 150-bp paired-end platform (BGI, China).

The *Sulfitobacter* dataset comprised 269 public genomes available in NCBI GenBank (on 14 July 2023) and 10 new genomes from strains closely related to *S. pontiacus* EE-36 (16S rRNA gene identity > 99.7%) and isolated from global ocean samples (Table S9).

Only four public genomes of *Dinoroseobacter* are available in NCBI GenBank. These strains were isolated from dinoflagellates (Table S9). Since two of the genomes, namely *Dinoroseobacter* shibae DSM 112351 and *Dinoroseobacter* shibae DFL-12 (= DSM 16493), were derived the same bacterial isolate, only the later was kept for downstream analysis. The remaining three strains shared high sequence similarity (> 98% ANI) and were assigned to a single population by PopCOGenT including three falling into a clonal complex (Fig. S5). To obtain more non-redundant genomes contributing to the genetic diversity of *D. shibae*, we sequenced the genomes of four new *Dinoroseobacter* genomes which are closely related to *D. shibae* DFL-12 (16S rRNA gene identity > 99.7%) and three strains (DFL-16, DFL-27, and DFL-38) were isolated from the same culture of *Alexandrium ostenfeldii* (Culture KO287) and strain LW35 was from the bacterial biofilm of the marine brown alga, *Fucus spiralis* (Table S9).

All new isolates of CHUG and *Dinoroseobacter* were sequenced using the DNBSEQ 150-bp paired-end sequencing (BGI, China) platform, and all new isolates of *Sulfitobacter* were sequenced using the Illumina HiSeq platform. We did not sequence additional *Ruegeria* strains, as this genus has a great number of public genomes available in NCBI GenBank.

For each of newly sequenced genomes, adaptors and low-quality bases of raw reads were trimmed by Trimmomatic v0.36 ^14^ with the parameters of ‘ILLUMINACLIP: 2:30:10:1:true SLIDINGWINDOW:4:15 MAXINFO:40:0.9 MINLEN:40’, and the quality of remaining reads was checked by FastQC v0.11.5 (<http://www.bioinformatics.babraham.ac.uk/projects/fastqc>). Next, genomes were assembled using SPAdes v3.9.1 ^24^ with high quality reads, and the quality of assemblies was determined with CheckM v1.0.7 ^25^. Genes were predicted with Prokka v1.11 ^12^.

#### Population delineation and validation of population boundaries

PopCOGenT ^26^ was used to delineate populations based on the genome sequences of the natural isolates. The rationale of PopCOGenT is based on the fact that recent homologous recombination leads to identical regions between genomes by erasing the single nucleotide polymorphisms (SNPs). As such, strains subjected to frequent recent gene transfers are expected to show an enrichment of identical genomic regions, whereas SNPs are accumulated between genomes lacking recent gene flow. By calculating the length distribution of identical sequences among all the possible pairs of whole genome alignments, PopCOGenT can measure the amount of recent gene transfers, an integration of transfer rate and length of transferred DNA. If the observed length distribution deviates from the null distribution where identical sequences can only be changed by mutations, recent gene transfer is detected. Genomes showing less than 0.0355% divergence would be collapsed into clonal complex due to insufficient genetic divergence to identify recent gene transfer. Genomes are clustered into a putative population based on the amount of recent gene flow, and strains from different clusters and thus different populations show significantly reduced amount of recent DNA transfers. Genomes within a clonal complex are likely redundant, thereby underestimating $\pi_{s}$ of a population. For this reason, only one genome from a clonal complex was used for $\pi_{s}$ estimation. In most cases, we considered populations with a minimum of three non-redundant members for $\pi_{s}$ estimation. For each lineage studied here, their bacterial members inhabit a range of ecological niches (Fig. S2-4 and Table S9). However, our primary objective is to investigate the mechanisms behind genome reduction in bacterioplankton from oligotrophic surface ocean environments. As such, we focused on populations whose members were predominantly sourced from surface seawater for $\pi_{s}$ estimation.

For CHUG, two populations identified by PopCOGenT are of interest: CHUG_MC0 consisting of 21 non-redundant strains and CHUG_MC1 containing four non-redundant strains (Fig. S2). Except for strain HKCCA0882 isolated from the brown algal tissue, all members in these two populations were isolated from the surface seawater (Fig. S2). The CHUG_MC0 forms a paraphyletic group, with a clonal complex consisting of five strains (HKCCA1006, HKCCA1085, HKCCA1083, HKCCA1084, HKCCA1089) embedded within the CHUG_MC0 members in the phylogenomic tree (Fig. S2A). The $\pi_{s}$ of CHUG_MC0 estimated with and without this clonal complex lineage were 0.043 and 0.042, respectively.

Using the 279 *Sulfitobacter* strains, we identified six populations, each consisting of at least three non-redundant strains (Table S9). Given that most of the population members of *Sulfitobacter*_MC2, *Sulfitobacter*_MC5, and *Sulfitobacter*_MC6 were primarily sourced from the surface seawater (Table S9 and Fig S2), these three populations were used for $\pi_{s}$ estimation. Within population *Sulfitobacter*_MC2, its members do not form a monophyletic group. Instead, there is one monophyletic group comprising four strains (*Sulfitobacter*_MC2_sub1) that is separated from the remaining three strains (*Sulfitobacter*_MC2_sub2) in the phylogenomic tree (Fig. S3). Interestingly, the last common ancestor (LCA) of *Sulfitobacter*_MC2_sub1 and *Sulfitobacter*_MC2_sub2 gave rise to another 24 strains, and all of these descendants form a larger monophyletic group, which we refer to as *Sulfitobacter*_Clade1 (Fig. S3). To validate the recent gene flow between the *Sulfitobacter*_MC2 members is more frequent than that between *Sulfitobacter*_MC2 and the other 24 strains, we conducted an analysis of the length of identical segments in pairwise genomes within *Sulfitobacter*_Clade1. Members within *Sulfitobacter*_MC2 share an average of 492 bp identical segments for each pair of genomes, which is approximately twice the length of identical segments shared between members of *Sulfitobacter*_MC2 and the other 24 strains in *Sulfitobacter*_Clade1. This finding supports that recent gene flow homogenized members within *Sulfitobacter*_MC2, even when they do not form a monophyletic group. However, a population is defined as a biological species, which is expected to form a monophyletic group ^26^, so $\pi_{s}$ value estimated based on *Sulfitobacter*_MC2 may be biased. We thus only consider the $\pi_{s}$ values derived from *Sulfitobacter*_MC5 and *Sulfitobacter*_MC6.

For *Ruegeria*, 19 populations each with at least three non-redundant members were identified by PopCOGenT from 353 public genomes (Table S9 and Fig. S4). However, the number of strains from surface ocean within each population ranges from zero to three, we thus kept eight populations (*Ruegeria*_MC0, *Ruegeria*_MC2, *Ruegeria*_MC3, *Ruegeria*_MC6, *Ruegeria*_MC12, *Ruegeria*_MC13, *Ruegeria*_MC16, and *Ruegeria*_MC19) that had at least one member originating from the surface ocean for $\pi_{s}$ estimation. Within each of three populations (*Ruegeria*_MC3, *Ruegeria*_MC6, and *Ruegeria*_MC12), the members do not form a monophyletic group (Fig. S4). To validate the recent gene flow within each population, we summarized the length of identical genomic segments shared by pairwise genomes using the same method as mentioned earlier. Similarly, we determined the LCA of each population and used all descendants derived from that LCA (i.e., the extended clade) to identify identical genomic regions between all possible pairs of genomes.

The genome pairs were divided into two categories: i) both genomes were from the target population, and ii) one genome was from the target population and the other was from other members of the extended clade. Members within *Ruegeria*_MC3, *Ruegeria*_MC6, and *Ruegeria*_MC12 share approximately 2,048, 330, and 567 identical segments, with an average length of 545 bp, 2,581 bp, and 204 bp for each pair of genomes, respectively. These identical regions are over 33 times, 377 times, and two times longer, respectively, than the identical segments shared by members between the target population and the remaining clade members. These results again support the idea that recent gene flow connects members within a population defined by PopCOGenT, even when they are not members of a monophyletic group. Since a population is expected to form a monophyletic group, we ignored these three populations when estimating $\pi_{s}$.

The seven *Dinoroseobacter* genomes formed one population with six strains comprising a new clonal complex (Fig. S5). Hence, only one population with two genomes were non-redundant and used for $\pi_{s}$ estimation.

#### Effective population size (N_e_) estimation for natural populations

For each species, $\pi_{s}$ was estimated for each identified population, and *N_e_* was calculated accordingly. If more than two populations were available, the median value was used as the *N_e_* of that species. Otherwise, the mean *N_e_* was used if only two populations were available. To estimate $\pi_{s}$, the first step is to identify orthologous gene families using OrthoFinder-2.2.1 ^27^ shared by the genomes within each population. These single-copy genes were each aligned at amino acid level using MAFFT v.7.464 ^28^ and subsequently imposed on nucleotide sequences. Next, fourfold degenerate sites were identified for each gene alignment using ‘get4foldSites’ available at https://github.com/brunonevado/get4foldSites, and $\pi_{s}$ was calculated based on the following formulas ^29^:

$h_{i}= \frac{n}{n-1}$ × (1- $\sum p^{2})$, (1)

$\pi_{s}= \frac{\sum_{i=1}^{S} h_{i}}{N}$, (2)

where *n* is the number of strains for a given population, *p* is the allele frequency of each nucleotide at a segregating fourfold degenerate site, *h_i_* is the average pairwise genetic distance at the *i*th segregating fourfold degenerate site, *S* is the number of segregating sites within all fourfold degenerate sites, and *N* is the number of all fourfold degenerate sites. Finally, the median $\pi_{s}$ across all single-copy gene families were used to calculate the *N_e_* for each population.

#### Construction of phylogenomic tree of Roseobacter lineages

The 74 representative roseobacters were collected from NCBI GenBank database (14 July 2023) by following previous studies ^1,30^. Then, 120 conserved single-copy genes (bac120) were extracted from each genome, aligned using GTDB-tk v1.7.0 ^31^, and concatenated. Next, the phylogenomic tree was constructed using IQ-TREE v2.2.0 ^32^ with parameter ‘-m LG+C20+F+G -g backbone_tree’ and visualized using iTOL ^33^.

#### Regression analysis between mutation rate, effective population size, and genome size

We assessed the pairwise linear relationship between mutation rate, effective population size (*N_e_*), and genome size among the four Roseobacter species using generalized linear model (GLM) and phylogenetic generalized least square (PGLS) methods, which were implemented in ‘stats’ ^34^ and ‘caper’ ^35^ packages in R. 4.2.3 ^34^, respectively. For the GLM regression, the outlier data point (Bonferroni *p* < 0.05) was identified using the ‘outlierTest’ function in ‘car’ package ^36^. The PGLS analysis accounts for potential autocorrelation among closely related species, which may lead to artificial scaling relationship between traits. Based on a given phylogeny, this analysis can evaluate evolutionary association between trait, indicated by Pagel’s λ ^37^, with λ values ranging from 0 (no phylogenetic signal) to 1 (strong phylogenetic signal due to Brownian motion). Here, the input phylogeny of the four Roseobacter species was pruned from the phylogenomic tree of Roseobacter lineages constructed above. Given the significant phylogenetic effect on the relationship between genome size and effective population size ($\lambda$ = 1), the GLM regression analysis between these two traits may not be reliable.

Another 31 phylogenetically diverse species were included to investigate the relationship between mutation rate, *N_e_*, and genome size across prokaryotic lineages. The mutation rates of these 31 species have been collected in our previous study, and 22 of them have *N_e_* estimated ^13^. We updated the *N_e_* value for *Salmonella enterica* based on the recent mutation rate measurement of this species by Pan et al. (2022), which is only ~1/6 of that previously reported ^39^. We applied the same GLM and PGLS methods for regression analysis on a total of 35 species. The species tree phylogeny, used as input for the analysis, was constructed for these 35 species based on our prior study ^13^.

### Supplemental Results

#### Mutation rate of Sulfitobacter pontiacus EE-36 and Dinoroseobacter shibae DFL-12

For *Sulfitobacter*, a total of 111 MA lines were initiated from a single ancestral colony of strain *S.* *pontiacus* EE-36, and 110 of which survived after 188 transfers with each line undergoing 4,102 cell divisions (corrected with death rate). Of the surviving lines, 58 MA lines yielded high quality reads (see Methods) and were thus used for mutation calling. In total 57 MA lines accumulated mutations, yielding a total of 284 BPSs, 22 deletions and 14 insertions (Table S3). Three genes showed significant enrichment of mutations, containing four BPS, four deletions, and three insertions (bootstrap test; *p* < 0.05 for each gene) (Fig. 1B), which may have inactivated the three genes either by nonsense mutation, deletion or insertion mutation (Table S4). After ignoring those four BPSs, the genomic base-substitution mutation rate (μ) was calculated as (3.61 ±1.86) × 10^-10^ base substitutions per site per cell division (95% CI 3.20 × 10^-10^ – 4.06 × 10^-10^). To further exclude the effect of epistatic interaction caused by enriched mutations, seven MA lines with mutations occurring in these three genes were ignored, and the remaining 51 lines yielded 243 BPSs, 11 insertions and 18 deletions (Table S3), leading to a μ of (3.57 ± 1.89) × 10^-10^ (95% CI: 3.13 × 10^-10^ – 4.03 × 10^-10^), which was not significantly different from the mutation rate (3.96 ± 1.58) × 10^-10^ (95% CI: 2.79 × 10^-10^ – 5.45 × 10^-10^) derived from the remaining seven MA lines mutated at aforementioned genes but with these mutations excluded from the calculation (Wilcoxon–Mann–Whitney test, *p* = 0.404). No selection was detected in the MA experiment of *S. pontiacus* EE-36, as the ratio of accumulated mutations in intergenic sites to those in protein-coding sites (36 vs 207) did not differ significantly with that of intergenic to protein-coding sites (306,118 vs 2,421,042) (χ^2^ test, *p* = 0.094), and the ratio of accumulated nonsynonymous to synonymous mutations (159 vs 48) showed no significant difference from that of nonsynonymous to synonymous sites (1,855,162 vs 565,880) (χ^2^ test, *p* = 1).

For *Dinoroseobacter*, a total of 220 MA lines were initiated from a single ancestral colony of *D. shibae* DFL-12, 187 of which survived after 75 transfers with each line undergoing 1,710 cell divisions (corrected with death rate). Of the 149 lines that showed good sequencing quality (see Methods), 128 lines accumulated mutations, yielding a total of 296 BPSs, 34 deletions and 82 insertions (Table S5). Eight genes showed significant enrichment of mutations, containing 78 BPS, one deletion, and 63 insertions (bootstrap test; *p* < 0.05 for each gene) (Fig. 1C). Mutations in these eight genes may have inactivated the eight genes either by nonsense mutation (63), deletion (one) or insertion mutation (63) (Table S6). After ignoring those 78 BPSs, the μ was calculated to be (2.05 ± 2.20) × 10^-10^ (95% CI: 1.80× 10^-10^ - 2.35 × 10^-10^). Besides, 87 MA lines with mutations occurring in these eight genes were ignored owing to the possible effect of epistatic interaction caused by enriched mutations, which yielded 80 BPS, seven insertions and nine deletions in the remaining 62 MA lines. This translates to a μ of (1.81 ± 2.25) × 10^-10^ (95% CI: 1.44 × 10^-10^ – 2.26 × 10^-10^) (Table S5). This mutation rate is not significantly different from the one (2.22 ± 2.15) × 10^-10^ (95% CI: 1.88 × 10^-10^ – 2.64 × 10^-10^) derived from the remaining 87 MA lines mutated at aforementioned genes but with these mutations excluded from the calculation (Wilcoxon–Mann–Whitney test, *p* = 0.127). The ratio of accumulated mutations in protein-coding sites to those in intergenic sites (60 vs 20) is significantly smaller than that of protein-coding sites to intergenic sites (3,924,027 vs 467,836) (χ^2^ test, *p* < 0.001), suggesting that protein-coding regions accumulated fewer mutations than expected. A similar pattern has also been observed in previous studies ^40–44^. Besides, the ratio of accumulated nonsynonymous to synonymous mutations (41 vs 19) is significantly smaller than that of nonsynonymous to synonymous sites (3,190,340 vs 733,687) (χ^2^ test, *p* = 0.016), indicating that nonsynonymous sites accumulated fewer mutations than expected. These patterns imply the influence of purifying selection in removing strongly deleterious nonsynonymous mutations. Furthermore, the ratio of nonsynonymous mutations to all other mutations occurred in synonymous and intergenic sites (41 vs 79) is significantly smaller than the expectation (3,190,340 vs 1,201,523) (χ^2^ test, *p* < 0.001), reinforcing the role of purifying selection in preventing the fixation of deleterious nonsynonymous mutations, which led to fewer mutations accumulated in protein-coding and nonsynonymous sites than expected.
